# Inhibition of BCR::ABL1 tyrosine kinase activity Aids in the Generation of Stable Chronic Myeloid Leukemia Induced Pluripotent Stem Cells

**DOI:** 10.1101/2023.06.01.543015

**Authors:** Esther Sathya Bama Benjamin, Dinesh Babu, Gaurav Joshi, Bharathi M Rajamani, Krittika Nandy, Sonam Rani, Senthamizhselvi Anandan, Chitra Premkumar, Madhavi Maddali, Aby Abraham, Shaji R Velayudhan, Poonkuzhali Balasubramanian

## Abstract

Induced pluripotent stem cells (iPSCs) generated from patients with chronic myeloid leukemia (CML) have the potential for disease modeling to study disease pathogenesis and screening therapeutic interventions. In this study, we aimed to generate iPSCs from CD34^+^ hematopoietic progenitors of CML patients with varying responses to tyrosine kinase inhibitor (TKI) therapy. The generated CML-CD34-iPSC colonies displayed atypical “dome-shaped” morphology and underwent spontaneous differentiation in a few days. However, supplementation with imatinib (IM), the most widely used TKI to treat CML patients, in the culture medium improved the stability and maintenance of all isolated CML-CD34-iPSC colonies, allowing them to be maintained for more than 20 passages without significant differentiation. In contrast to previous studies, our results indicate that suppressing the BCR::ABL1 oncogenic pathway is essential for efficiently generating stable CML-iPSC colonies. Furthermore, we successfully differentiated these iPSCs to CD34^+^ hematopoietic progenitors both in the presence and absence of IM. This robust protocol for generating CML-iPSCs provides a valuable resource for disease modelling. The generated iPSCs will be a valuable tool for investigating CML pathophysiology, drug resistance mechanisms, and drug screening to identify novel and effective therapies for this disease.

## Introduction

Chronic myeloid leukemia (CML) is a myeloproliferative neoplasm that arises from hematopoietic stem cells (HSCs) transformed by BCR::ABL1 oncogenic fusion protein with constitutive serine/threonine kinase activity expressed in these cells. BCR::ABL1 triggers various downstream signaling pathways, including RAS/RAF/MEK/ERK, JAK2/STAT, and PI3K/AKT/mTOR pathways, leading to the accumulation of granulocytes at various stages of myeloid maturation (1,2). Although targeted therapy with tyrosine kinase inhibitors (TKIs), such as imatinib (IM), dasatinib (DA), and nilotinib (NIL), has dramatically improved the long-term survival of CML patients, only around 50% of patients achieve treatment-free remission (TFR) (3). The persistence of leukemic stem cells (LSCs) is one of the primary causes of disease recurrence (4). Due to the rarity of the LSC population, it is challenging to obtain sufficient cells for research to understand the mechanisms of TKI resistance of these cells and screen drugs to eradicate them (4,5).

Induced pluripotent stem cells (iPSCs) derived from patients with genetic diseases or cells with genetic alterations can be differentiated into scalable quantities of disease-relevant differentiated cells. Thus, they are a powerful tool for studying disease pathogenesis and drug screening (5). iPSCs generated from patients with leukemias help understand how oncogenes and patient-specific chromosomal abnormalities influence the development of leukemia-like phenotypes *in-vitro* (6). CML-iPSCs generated from the same patient at various stages of the disease and from patients with different responses to TKI therapy are valuable resources for investigating molecular and epigenetic mechanisms of CML progression and drug resistance, leading to the identification of novel therapeutic targets and drug candidates for the treatment of CML (7,8). Moreover, iPSCs can be efficiently genetically manipulated by gene editing methods to identify specific genes involved in disease pathogenesis (9).

Previous studies have demonstrated the successful generation of CML-iPSCs from peripheral blood mononuclear cells (10) and CD34^+^ hematopoietic stem and progenitor cells (HSPCs) isolated from CML patients (7,11–13). CML-iPSCs exhibit resistance to IM like CML-LSCs, so they are considered suitable for investigating the cellular mechanisms underlying IM resistance (12,14,15). CML-iPSCs could be differentiated into hematopoietic progenitors that recapitulated some of the pathophysiologic features of the disease (7,10,12–14). However, it remains unclear how *BCR::ABL1* expression and TKI treatment impact the efficiency and quality of iPSC generation from CML cells.

In this study, we describe the generation of iPSCs from the CD34^+^ cells of CML patients who exhibited varying responses to TKI therapy for disease modeling and investigating the effect of BCR::ABL1 oncoprotein expression on the maintenance of CML-iPSCs. In contrast to previous studies, we found that the expression and activity of BCR::ABL1 affect the maintenance of CML-CD34-iPSCs, and it was essential to suppress the TKI activity of BCR::ABL1 fusion protein by TKI supplementation for their survival and maintenance. The CML-iPSCs generated using our protocol could be differentiated into BCR::ABL1 positive hematopoietic progenitors. The patient-specific CML-iPSCs and the differentiated hematopoietic cells generated by this method have a wide range of applications, including high throughput CRISPR-Cas9 screening, to identify novel pathways involved in CML pathogenesis and drug resistance.

## Materials and methods

### Ethical statement

This study was approved by the institutional review board and the institutional bio-safety committee of Christian Medical College, Vellore.

### Isolation and expansion of CD34^+^ cells

Peripheral blood samples were obtained from a healthy donor (N=1), who underwent mobilization of CD34^+^ cells for hematopoietic transplantation, and newly diagnosed chronic phase-CML (CP-CML) patients (N=4) after obtaining written informed consents. Peripheral blood mononuclear cells (PBMNCs) were isolated using Ficol density gradient centrifugation. The CD34^+^ cells were purified from PBMNCs by the magnetic separation method (EasySep, Stem Cell Technologies) and were cryopreserved. The cryopreserved CD34^+^ cells of the CML patients and the normal donor were thawed and cultured in CD34^+^ cell expansion medium constituting StemSpan™ SFEM II medium (Stem Cell Technologies) supplemented with CD34^+^ Expansion Supplement and UM729 (Stem Cell Technologies). The cells were cultured for 6-7 days with half medium change every alternative day to assess the cell proliferation.

### Reprogramming of CML and normal CD34^+^ cells

The CD34^+^ cells were cultured in the expansion medium for 3-days and then nucleofected with episomal reprogramming plasmids (pCXLE-hOCT3/4-shp53 (Addgene: 27077), pCXLE-hSK (Addgene 27078) and pCXLE-hUL (Addgene: 27080) (kind gift from Shinya Yamanaka), as described previously (16). Electroporation was carried out using Neon Transfection System (ThermoFisher) at 1600V, 10ms, and 3 pulses. After five days of culture, the nucleofected cells were transferred to Matrigel-coated plates containing Essential-8 (E8) medium (ThermoFisher Scientific). Individual colonies were hand-picked between day 21 and day 32 and were passaged and maintained in mTeSR Plus medium (StemCell Technologies) supplemented with 10µM IM (Sigma Aldrich) to generate stable iPSC clones.

### Western Blot

The iPSCs were dissociated using 1mM EDTA (ThermoFisher Scientific), and the protein lysates were prepared in radioimmunoprecipitation assay buffer supplemented with a protease inhibitor mixture (Roche Applied Science, IN, USA) and 2mM phenylmethylsulfonyl fluoride (Sigma-Aldrich). About 30μg of whole cell lysate was loaded in 10% SDS-polyacrylamide gel, and the proteins were transferred to a polyvinylidene difluoride membrane. The membrane was blocked with 10% non-fat dry milk powder in tris-buffered saline containing 0.1% Tween 20 (TBST) for two hours and then incubated with an anti-phosphoCRKL antibody (Cell Signaling Technology) and Anti β –Actin antibody (ThermoFisher Scientific). The proteins were detected using the ECL chemiluminescence kit (Thermo Scientific, Pierce), and the images were captured using the FluorChem E system (Protein Simple).

### Analysis of *BCR::ABL1* fusion transcripts

RNA was extracted from patients’ PBMNCs and iPSCs using TRI Reagent (Sigma-Aldrich). cDNAs were prepared from 2 μg RNA with random hexamers using the High-Capacity cDNA Synthesis Kit (Thermo Scientific), followed by RT-PCR to identify the *BCR::ABL1* fusion transcripts using a previously described protocol (17). The *BCR::ABL1* mRNA copies were assessed using an in-house ddPCR assay (Datari et. al., manuscript under preparation). iPSC colonies were screened for BCR::ABL1 kinase domain mutations using a previously described protocol (18).

### Alkaline phosphatase staining

Alkaline Phosphatase staining was performed using the Alkaline Phosphatase Detection Kit (Sigma-Aldrich) following the manufacturer’s instructions.

### Flow cytometry

To analyze the expression of TRA-1-60 in iPSCs, the colonies were dissociated using TrypLE (Thermo Fisher Scientific) and then stained using a PE-conjugated TRA-1-60 antibody (Thermo Fisher Scientific) for 20 mins. The cells were washed with mTeSR Plus (Stem Cell Technologies) containing Revitacell (Thermo Fisher Scientific), and the expression was analyzed using Navios (Beckman Coulter) or FACS Aria (BD Biosciences) flow cytometers. To measure the purity of CD34+ cells after their magnetic separation from PBMNCs, the cells were stained with an APC-conjugated anti-CD34 antibody (BioLegend).

### Immunofluorescence

Immunofluorescence for pluripotency markers, such as SOX2, TRA-1-81, OCT4A, SSEA-4, NANOG and TRA-1-60, was performed as previously described (19).

### Trilineage differentiation

Trilineage differentiation of iPSCs to ectoderm, endoderm and mesoderm lineages was carried out using STEMdiff tri-lineage differentiation Kit (Stem Cell Technologies) as per the manufacturer’s protocols.

### Hematopoietic differentiation of iPSCs

For the hematopoietic differentiation of iPSCs, we used the STEMdiff™ Hematopoietic Kit (Stem Cell Technologies). Briefly, iPSCs were either single-cell sorted or seeded as small aggregates in mTeSR Plus (Stem Cell Technologies). Subsequently, the culture medium was changed to STEMdiff™ A medium, with or without IM. After three days, 70,000 cells were reseeded in STEMdiff™ A medium and maintained for another three days. Following this, the medium was switched to STEMdiff™ B. After 12 days of culture, the cells present in suspension were harvested and analyzed by flow cytometry for the expression of hematopoietic surface markers.

### Colony-Forming Assay

In each experiment, 7,000 hematopoietic progenitors generated from iPSCs were plated onto MethoCult GF H4636 medium (Stem Cell Technologies). The number of colonies in each experiment was scored on day 14 following the manufacturer’s instructions.

## Results

### Generation of iPSCs from normal and CML CD34^+^ cells

As CD34^+^ cells of CML patients are known to constitute the leukemia stem/progenitor cells (LSPCs) that are associated with disease recurrence (4,20), we aimed to reprogram CML-CD34^+^ cells from two patients who achieved complete cytogenetic response and one who achieved suboptimal response (>1% *BCR::ABL1* transcripts detected in peripheral blood cells after 12months) after TKI therapy (**Suppl Table 1**) and a healthy volunteer as described (**Figure 1A**). CD34^+^ cells purified from CML patients and those from a healthy donor were cultured in CD34^+^ expansion medium. The number of CD34^+^ cells increased by 15 to 20-fold (Mean: 20.2±:3.34) without reducing the percentage of CD34^+^ cells (**Figure S1A** and **S1B**).

**Figure 1.**
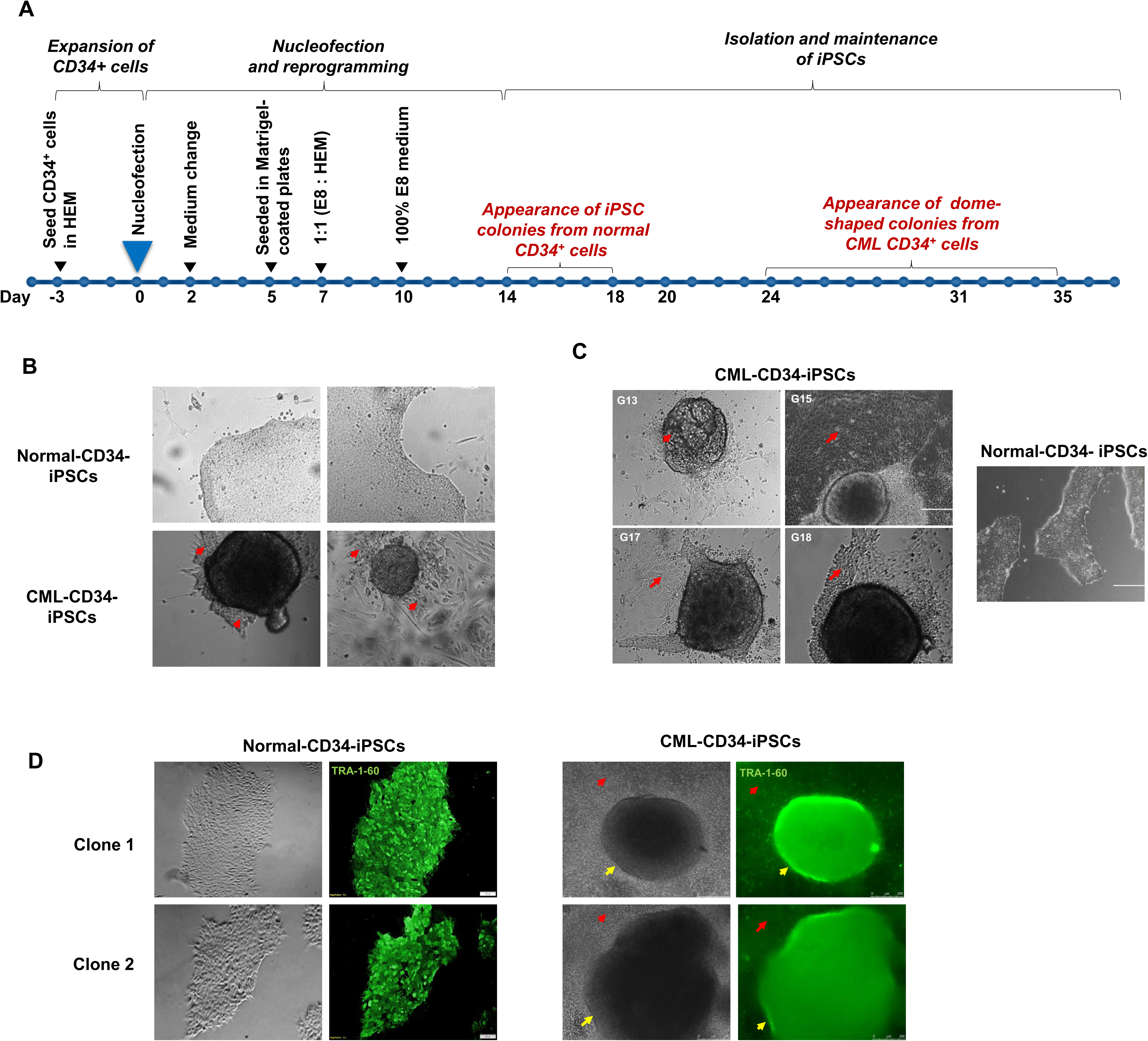

We nucleofected proliferating CD34^+^ cells from day 3 of the culture with episomal reprogramming plasmids (13). The normal CD34^+^ cells gave rise to iPSC colonies with typical morphology between day 14 and day 18 (**Figure 1B**). These colonies were hand-picked between day 20 and day 24 to generate stable iPSC clones. The normal iPSC clones maintained typical iPSC morphology without differentiation for 12 passages and expressed high levels of pluripotency markers (**Figures 1C** and **1D**). In contrast, CD34^+^ cells from TKI-responsive CML patients did not form typical iPSCs but formed several “dome-shaped” colonies after 30 days (**Figures 1B and C**). These colonies were surrounded by adherent cells (**Figure 1C)** that resembled those formed by the spontaneous differentiation of iPSCs. We isolated 11 “dome-shaped” colonies from 3 patients and cultured them for 4 to 5 passages. They also expressed TRA-1-60 in the initial passages (**Figure 1D**). During subsequent passages, the “dome-shaped” colonies peeled off from the bottom of the plates and floated in the medium, leaving the differentiated cells attached to the plate.

### IM supplementation aids in the maintenance, stability, and pluripotency of CML iPSCs

Previous studies showed that CML-iPSCs express *BCR::ABL1* transcripts present in the donor cells used for reprogramming (10–12,14). In our study, the “dome-shaped” CML-CD34-iPSC colonies also expressed the same types of *BCR*:*:ABL*1 fusion transcripts (e13a2 and e14a2) as those present in the respective patients’ peripheral blood cells used for reprogramming (**Figure S2A** and **S2B**). Additionally, karyotyping confirmed the presence of t(9;22) in all iPSC clones (**Figure S2C**), and none of the clones had mutations in the *BCR::ABL1* kinase domain.

As it has been reported that TKI treatment enhances pluripotency marker expression in BCR::ABL1+ iPSC clones (15), we hypothesized that the atypical morphology and high rate of spontaneous differentiation observed in our CML-CD34-iPSC colonies were due to the constitutive expression of BCR::ABL1 fusion protein and its tyrosine kinase (TK) activity. To inhibit the TK activity of the BCR::ABL1 fusion protein, we supplemented the medium with 10µM IM and monitored the colonies for changes in their morphology characteristics. We observed that within two days of IM supplementation, the colonies gained the typical iPSC morphology (**Figure 2A**) with an increase in the TRA-1-60+ cell population (**Figures 2B** and **2C**). All the clones were positive for pluripotency marker expression (**Figure 2D**). They also could be differentiated into three germ layers, ectoderm, endoderm, and mesoderm (**Figure S3**). When IM was withdrawn, the IM-supplemented CML-CD34-iPSC colonies reverted to the “dome-shaped” morphology within 2-4 days (**Figure 3A**), with a reduction in cell viability (**Figure 3B**) and TRA-1-60 expression (**Figure 3C).** In contrast, when the control iPSCs were treated with IM, there was a reduction in the colony number, suggesting that IM supplementation specifically enhances the maintenance of CML-CD34-iPSCs (**Figure S4**).

**Figure 2.**
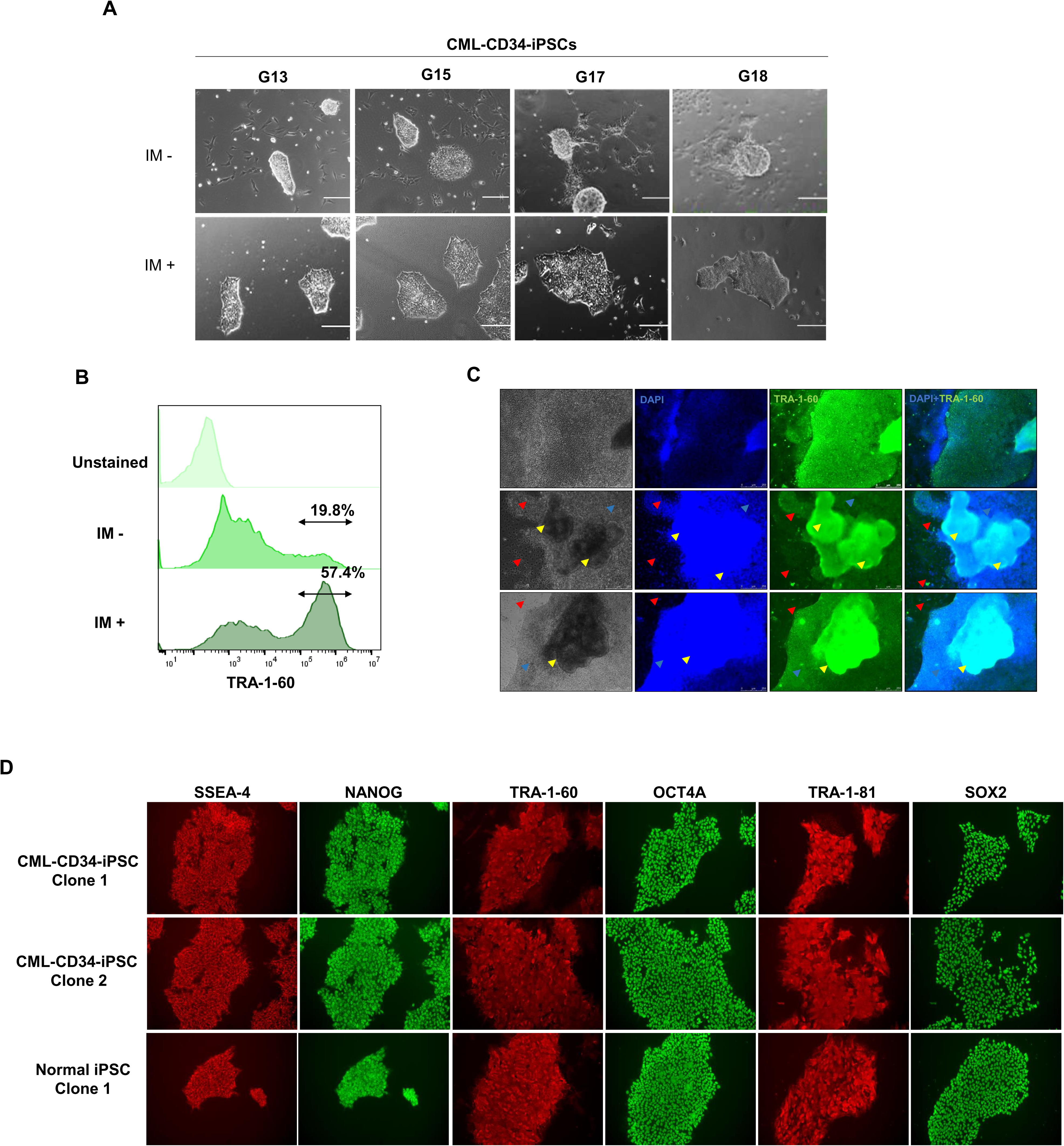

**Figure 3.**
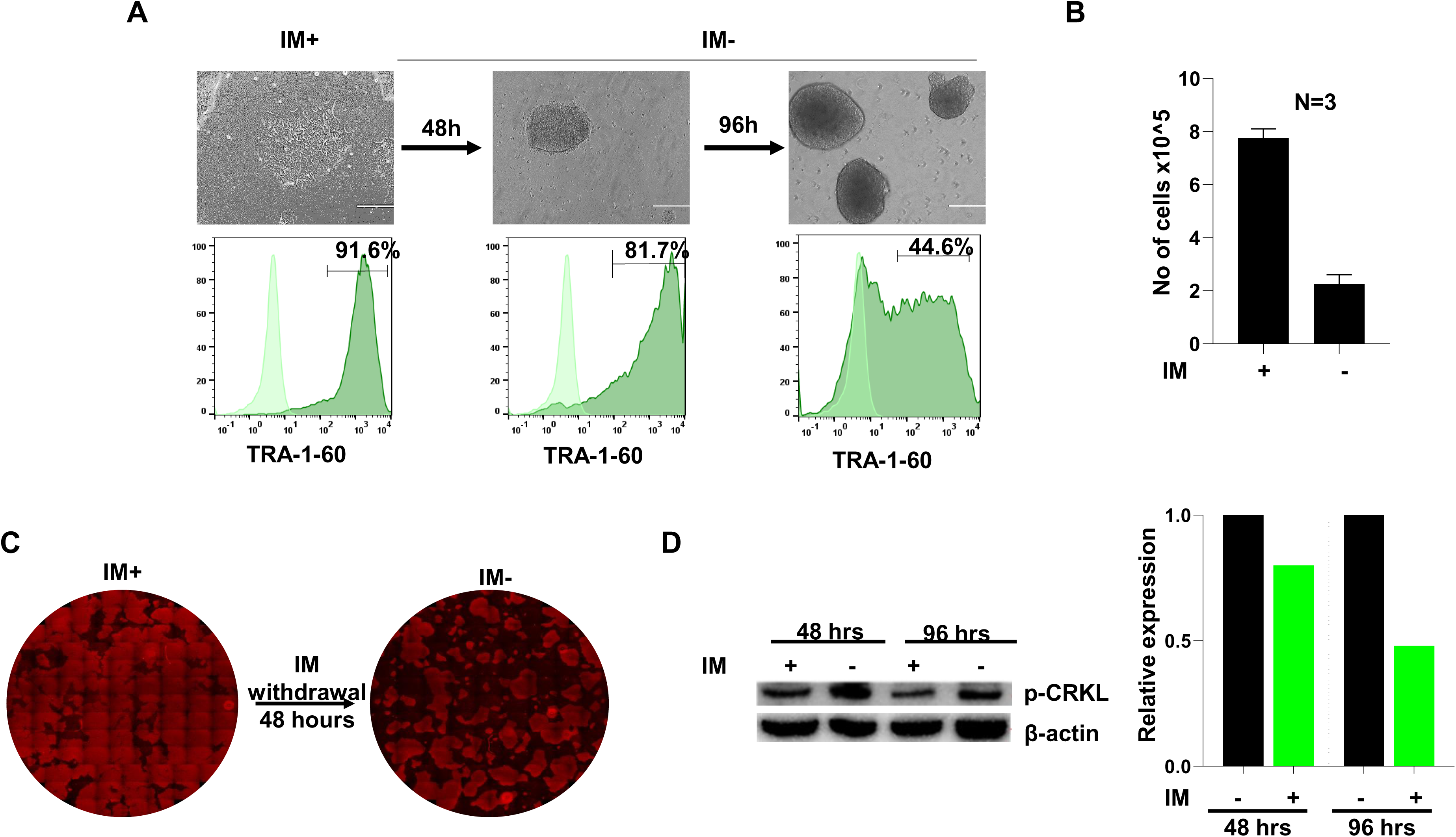

CRK-like proto-oncogene (CRKL) is a downstream target of BCR::ABL1 and is phosphorylated by BCR::ABL1 fusion protein (20,21). As p-CRKL levels can be used to determine the status of BCR::ABL1 kinase activity, the expression of p-CRKL was analyzed in CML-CD34-iPSCs before and after the IM treatment (**Figure 3D**). It was found that p-CRKL expression was decreased in CML-CD34-iPSCs after IM treatment, suggesting that suppression of BCR::ABL1 mediated tyrosine phosphorylation resulted in the stable maintenance of CML-CD34-iPSCs.

### The effect of IM in reprogramming of CML-CD34+ cells

As IM supplementation inhibits BCR::ABL1 kinase activity and enhances the stability and pluripotency of CML-iPSCs, we investigated the effect of IM on the reprogramming of CML-CD34^+^ cells. After nucleofection of CD34+ cells from two IM-responsive patients, the cells were cultured in the reprogramming medium supplemented with 10µM IM (**Figure 4A**). Reprogramming without IM supplementation generated “dome-shaped” colonies after 3 weeks, and they gained typical iPSC morphology after IM supplementation with increased TRA-1-60 expression (**Figure 4B**). However, continuous IM supplementation throughout reprogramming did not generate “dome-shaped” colonies or colonies with typical morphology. These findings suggest that while IM enhances the stability of CML-iPSCs by inhibiting *BCR::ABL1* kinase activity, it impedes the reprogramming of CML-CD34^+^ cells. Overall, our results demonstrate that IM supplementation transforms the “dome-shaped” colonies formed during the reprogramming of CML-CD34^+^ cells into iPSCs with typical flat morphology. Furthermore, IM improves the pluripotency of CML-CD34-iPSCs by inhibiting the BCR::ABL1-mediated TK activity.

**Figure 4.**
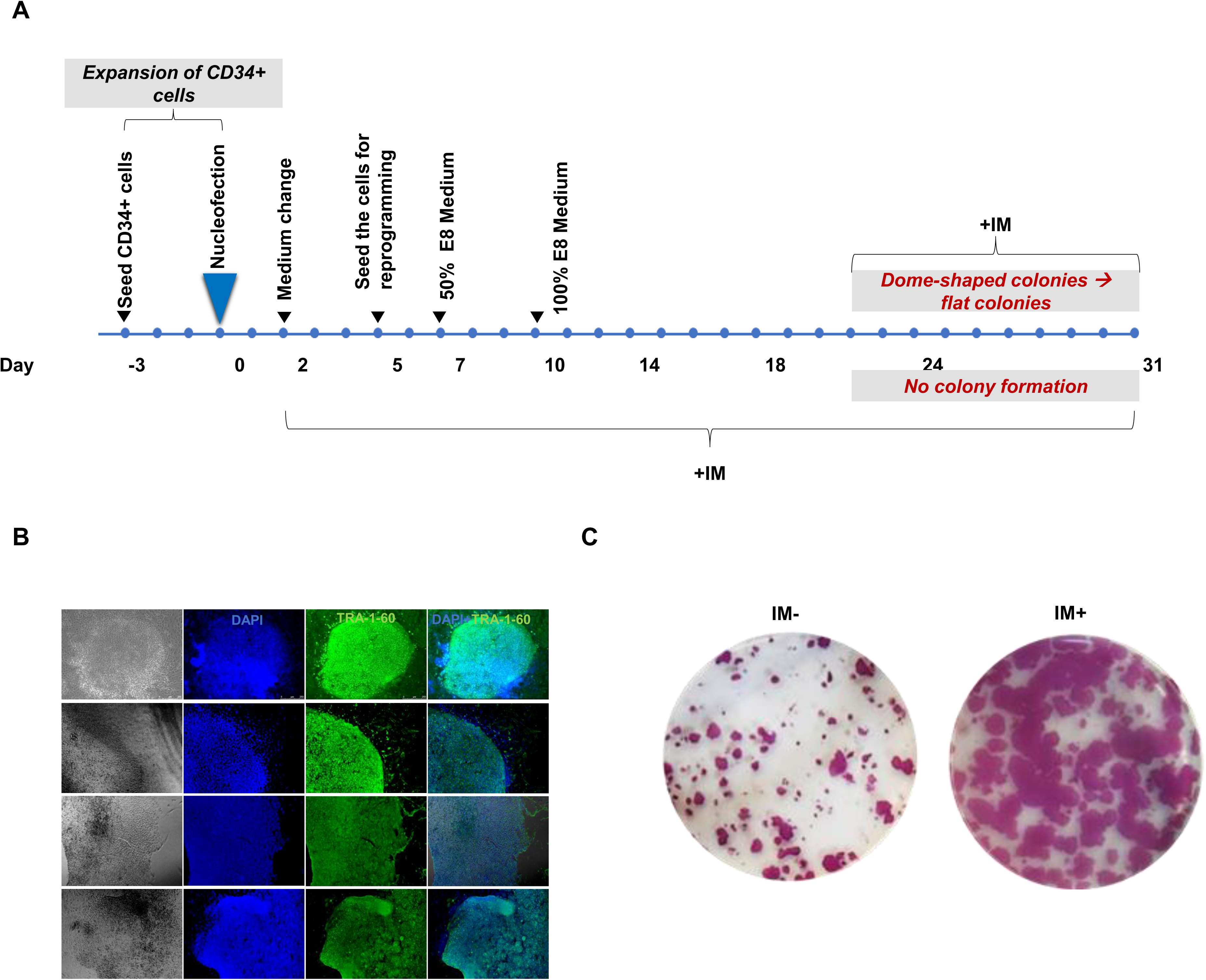

### Reprogramming of CD34^+^ cells from an IM refractory CML patient

The HPCs derived from the iPSCs generated from IM refractory CML patients offer a valuable tool to investigate the mechanisms underlying IM therapy resistance and develop a more effective treatment for those who do not respond well to IM. Therefore, we reprogrammed CD34^+^ cells from an IM refractory patient, with and without IM supplementation, during the reprogramming process (**Figure 4A**). In contrast to our observation in the IM responders’ cells, we observed an early emergence of iPSC colonies from the IM-refractory patient’s cells (∼3 weeks Vs. ∼4 weeks). However, reprogramming of the cells from the IM refractory patient-generated more colonies with continuous IM supplementation than without IM (**Figure 4C**). We isolated 9 colonies, 8 of which exhibited “dome-shaped” morphology (**Figure 1C**), while 1 showed typical iPSC morphology (**Figure S5A**). Karyotyping analysis of the “dome-shaped” colonies generated from the IM refractory patient demonstrated the presence of t(9;22), whereas the one clone with the typical iPSC morphology showed a normal karyotype (**Figure S5A** and **S5B**). These results suggest that the “dome-shaped” morphology of the CML-CD34-iPSCs is because of BCR::ABL1 expression in these cells.

### Second and third generation TKIs also aid in the maintenance of CML iPSCs

Second generation TKIs, DA and NIL, and third generation TKI ponatinib were approved for patients who develop resistance to or are intolerant to first or second-generation TKIs, respectively. These TKIs target specific mutations in the BCR::ABL1 fusion protein, providing more effective and durable responses in CML patients. We explored the effects of second and third generation TKIs, DA, NIL, and ponatinib, in the maintenance of CML-CD34-iPSCs. We replaced IM with different concentrations of DA, NIL, and ponatinib and evaluated their effects on CML-CD34-iPSCs. CML-CD34-iPSCs treated with these TKIs generated stable the CML-CD34-iPSCs with 2nM DA, 40nM NIL, and 2nM ponatinib, while at lower concentrations, resulted in “dome-shaped” colonies (**Figure 5**). Together, our results suggest that the inhibition of BCR::ABL1 TK activity promotes the pluripotency of CML-iPSCs.

**Figure 5.**
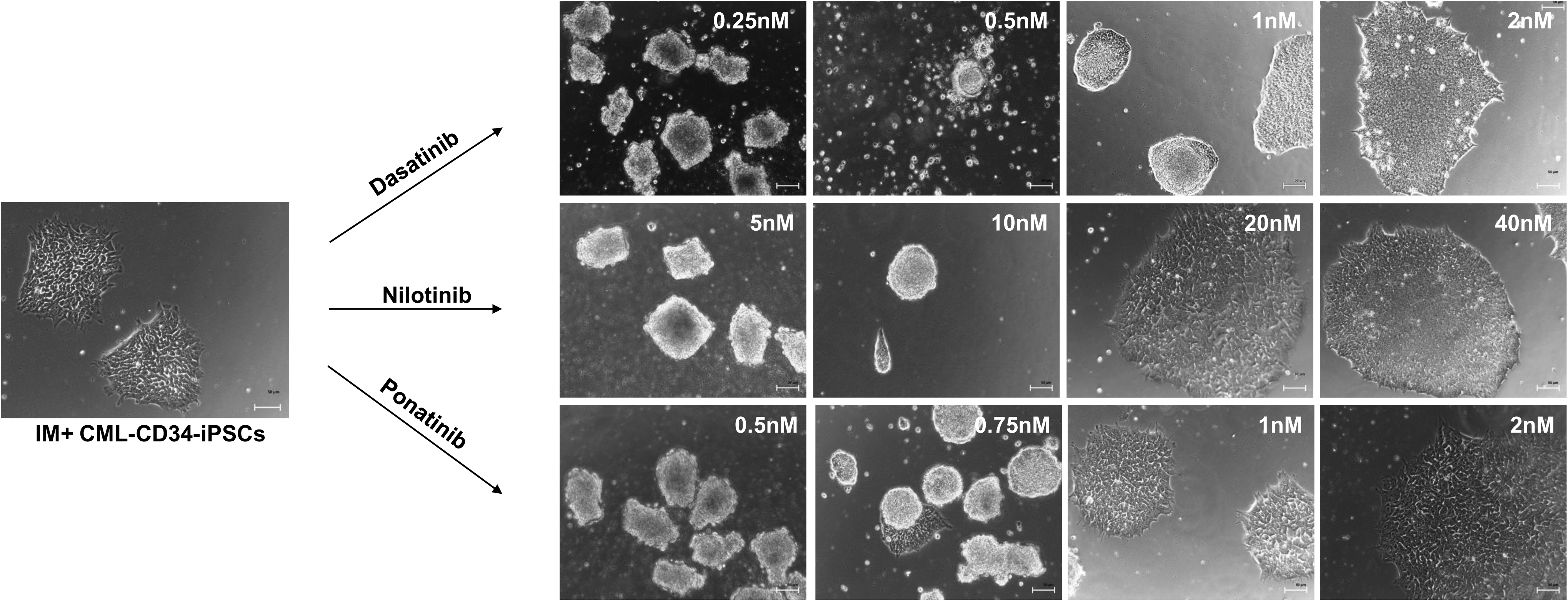

### Hematopoietic differentiation of CML iPSCs

Previous studies have demonstrated that CML-iPSCs can be successfully differentiated into hematopoietic progenitor cells (HPCs), despite the expression of BCR::ABL1 oncoprotein and its TK activity in these cells (7,14,22). However, the efficiency of hematopoietic differentiation of CML-iPSCs has been reported to be lower than normal iPSCs (7,10), and the resulting HPCs exhibited a partial recovery of TKI sensitivity (13). Considering our finding that IM supplementation is necessary for the maintenance of CML-CD34-iPSCs, we investigated whether IM supplementation is also required for the differentiation of CML-iPSCs into HPCs.

To address this question, we initially performed single-cell sorting to obtain iPSC colonies with an equal number of cells, which were subsequently subjected to hematopoietic differentiation (**Figures 6A and B**). We incorporated IM supplementation at different stages of the differentiation process (**Figure 6B**). We differentiated CML-CD34-iPSCs using two different concentrations of IM (1µM and 2µM) as well as without any IM supplementation. We could obtain suspension cells constituting of HPCs both in the presence and absence of IM (**Figure S6**). Interestingly, CML-CD34-iPSCs differentiated without IM before and after mesoderm induction (E1) (**Figure 6B**) showed a 5-fold increase in the yield of CD34^+^CD45^+^ HPCs and the total number of suspension cells compared to those differentiated in the presence of IM throughout the entire differentiation (E3 and E5) (**Figure 6D**). Flow cytometry analysis of hematopoietic markers revealed that approximately 41% of the suspension cells were CD34^+^CD45^+^ HPCs (**Figure 6C**). Utilizing single-cell sorting of iPSCs, we were able to generate a sufficient number of HPCs for a drug screening experiment.

**Figure 6.**
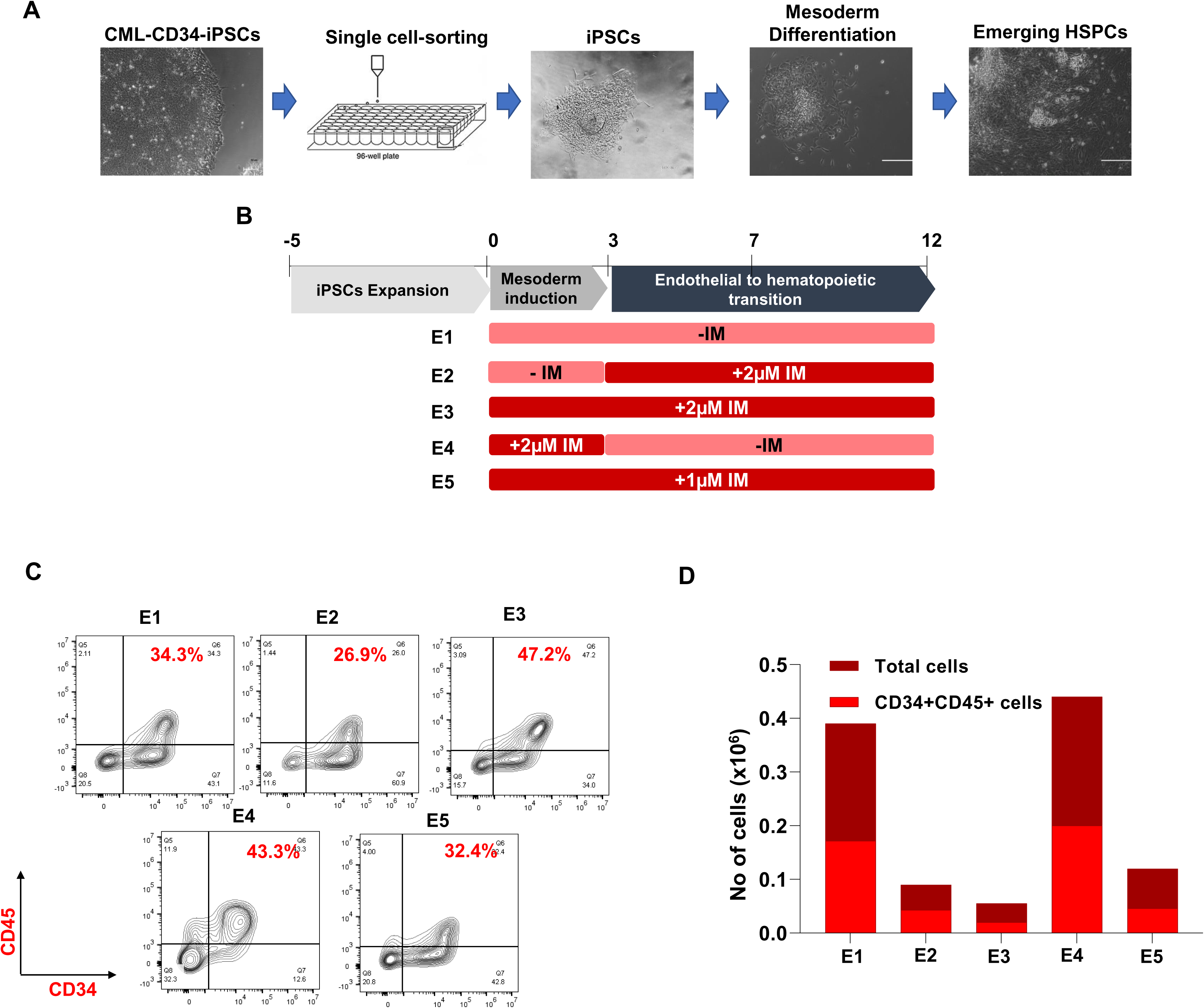

Based on the results of this experiment, we seeded iPSCs as small aggregates and then subjected differentiation with and without IM supplementation after mesoderm induction (**Figure 7A)**. Notably, the CML-CD34-iPSCs differentiated without IM yielded approximately 30 times more HPCs **(Figure 7C**) compared to differentiation with IM supplementation. The decline in HPC differentiation observed in the presence of IM can be attributed to the sensitivity of CML-iPSC-HPCs to IM. Flow cytometry analysis revealed that around 50% of the suspension cells constituted the CD34^+^CD45^+^ population of HSPCs (**Figure 7B)**. Furthermore, hematopoietic colony formation showed that the HPCs generated without IM had more myeloid progenitors compared to those derived from a normal iPSC clone (**Figure 7D**). To further validate these findings, we differentiated CML-CD34-iPSCs derived from two additional patients **(Figure 7E**). Flow cytometry analysis showed 40% to 70% CD34^+^CD45^+^ cells in the absence of IM from these two CML-CD34-iPSCs (**Figure 7G)**. Consistently, it was observed that differentiation in the absence of IM resulted in more HPCs compared to the other experimental groups (**Figure 7F**).

**Figure 7.**
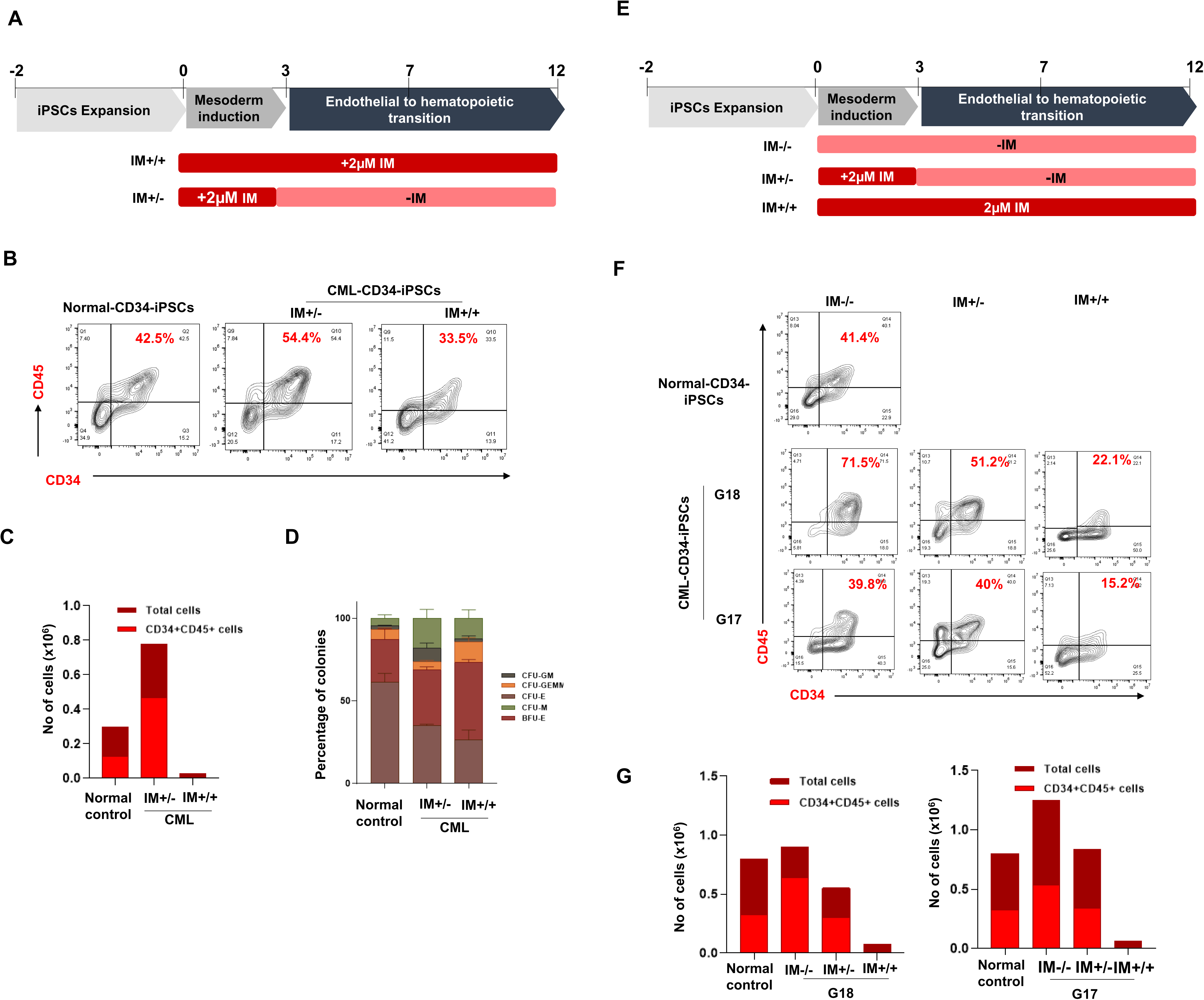

## Discussion

Eliminating CML-LSCs is a critical objective in the treatment of CML, as it can prevent disease recurrence, extend survival, improve quality of life, and enable curative treatments (3,4,23). However, the rarity of these cells poses a challenge in studying the molecular mechanisms that contribute to their survival. Generating iPSCs from CML CD34^+^ cells can help overcome this hurdle. These cells can be expanded and redifferentiated into hematopoietic cells, providing a continuous supply of cells for extensive proteome, epigenome, transcriptome, and drug screening studies.

Several groups have generated CML iPSCs from PBMNCs (10) and CD34^+^ cells (7,11,12,14) using lentiviral (14,15), retroviral (12,13), Sendai (11), or episomal vectors (7,10). However, these studies did not address the atypical morphology and low reprogramming efficiency of CML iPSCs. Additionally, the effect of BCR::ABL expression on the morphology and stabilization of CML-iPSC colonies (9) was not investigated.

We observed that the reprogramming efficiency of CML CD34^+^ was approximately 50 times lower than the normal CD34^+^ cells, as previously reported (7,11). One possible explanation for this low efficiency could be the presence of the BCR::ABL1 oncogene, which may disrupt the reprogramming process of CML CD34^+^ cells (24). Furthermore, CMLCD34^+^ cells formed colonies later than normal CD34^+^ cells, and they exhibited atypical colony morphology (14). Bedel et. al. reported that CML iPSCs had a distinct morphology resembling that of cell aggregates similar to mouse embryonic stem cells (12,14).

The instability and differentiation-prone nature of CML-CD34-iPSCs may be due to the presence of *BCR::ABL1* oncogene, which can interfere with the reprogramming factors required to generate stable iPSC colonies. Ito *et al*. demonstrated that the use of molecularly targeted drugs such as IM, which can suppress the activity of the BCR-ABL oncogene, can improve the transcriptional regulation of reprogramming factors and enhance pluripotency marker expression (24). This study also demonstrated that the expression of reprogramming factors in the BCR::ABL+ K562 cells increased pluripotency marker expression after IM supplementation, confirming that oncogene suppression improves the reprogramming of cancer cells. Our study confirmed these findings and showed that IM supplementation was necessary to generate stable CML iPSCs without spontaneous differentiation. IM supplementation improved the stability and maintenance of all isolated CML-CD34-iPSC colonies, allowing them to be cultured for over 20 passages without significant differentiation. The CML-CD34-iPSCs cultured with IM showed increased proliferation, while IM withdrawal decreased the proliferation and TRA-1-60 expression. These results were consistent with a previous study that reported faster growth of CML-iPSCs in the presence of TKI (14,15).

The phosphorylation of CRKL is decreased in IM-treated primary CD34^+^ HPCs isolated from CML patients (25) and CML iPSCs (**Figure 3D**), suggesting that the BCR-ABL TK activity in hematopoietic cells and CML iPSCs may involve similar pathways. The fact that IM could stabilize the pluripotency of CML-iPSCs uncovers a novel mechanism of pluripotency maintenance. Both CML-iPSCs and LSCs express BCR-ABL fusion protein but are IM resistant. Therefore, CML-iPSCs created using our strategy and the HPCs obtained from these iPSCs can be used for CRISPR-Cas9 screening to study the mechanisms of TKI resistance exhibited by these cells and discover drugs that can effectively eliminate CML-LSCs.

CML-iPSCs generated in the presence of IM could be successfully differentiated into HPCs. The iPSC-derived HPCs obtained from multiple patients will help investigate the survival mechanisms of BCR-ABL+ hematopoietic cells in TKI-resistant patients.

## Acknowledgments

We acknowledge the support of the core imaging and flow cytometry facilities at the Centre for Stem Cell Research (a unit of inStem, Bengaluru, Christian Medical College Campus, Bagayam, Vellore). We thank Ms. Sangeetha for her help in collecting and processing primary CML samples. We thank our clinical colleagues in the Department of Haematology for managing the patients with CML and providing the data on response to therapy.

---

**Figure S1.**
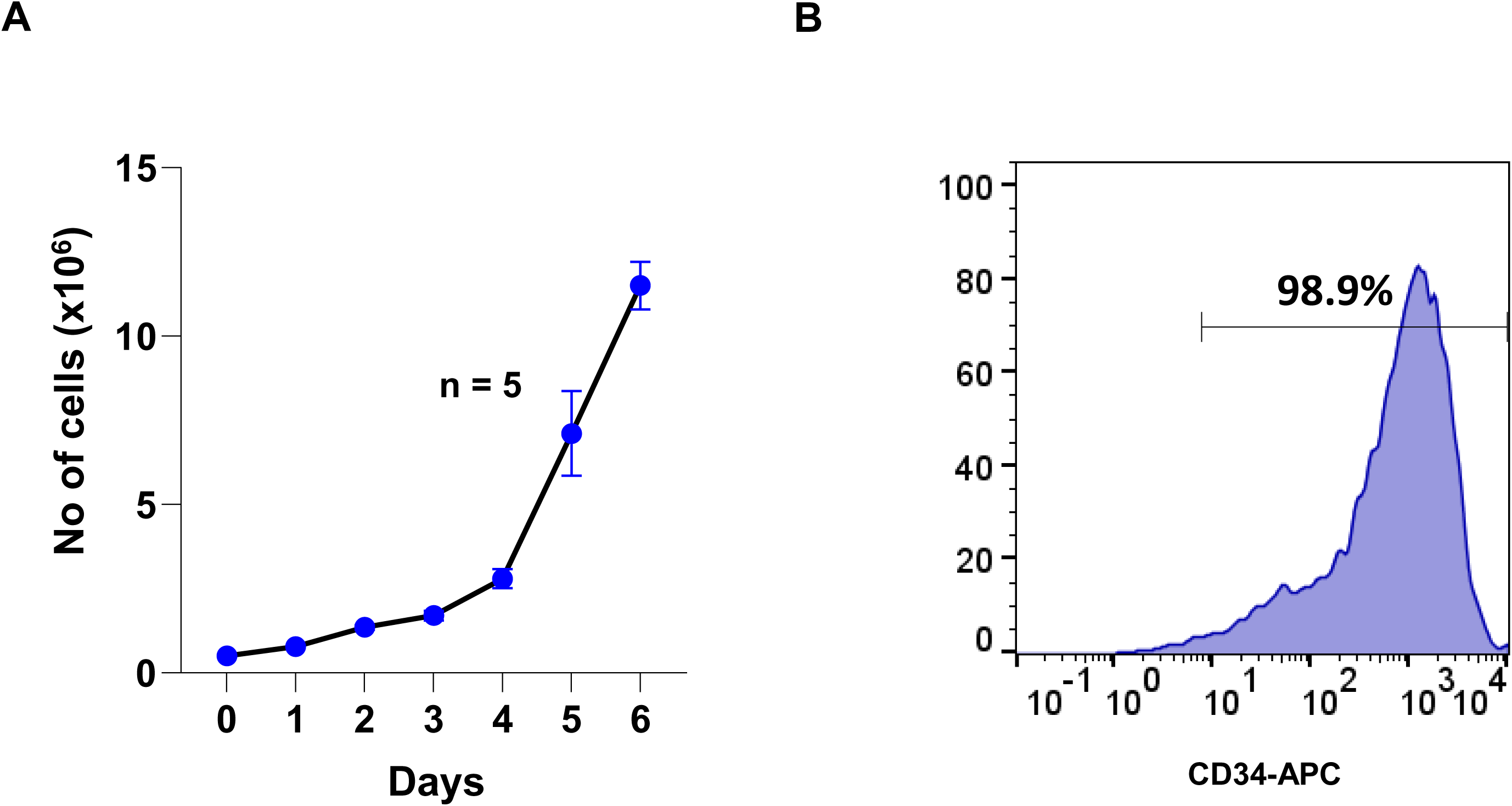

**Figure S2.**
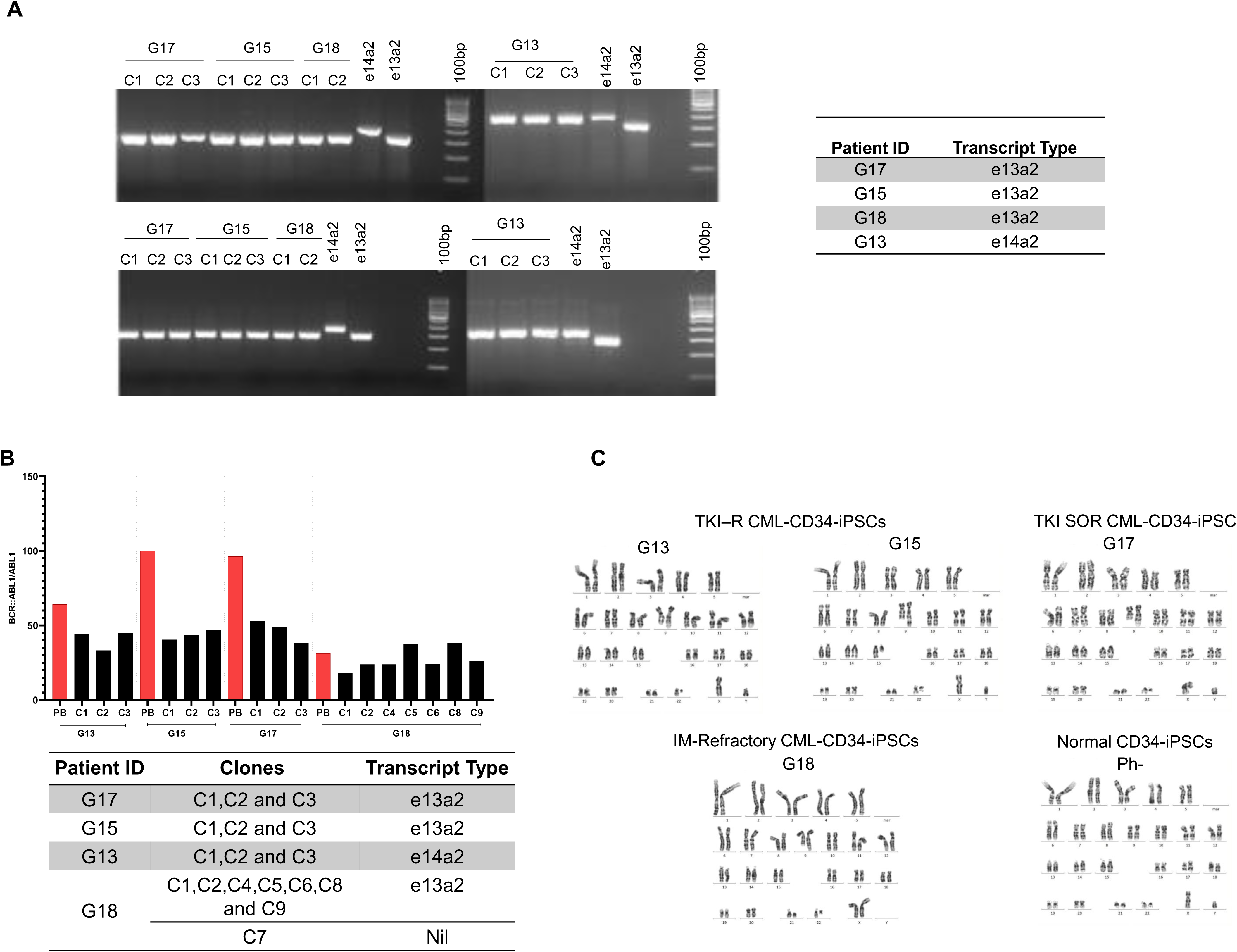

**Figure S3.**
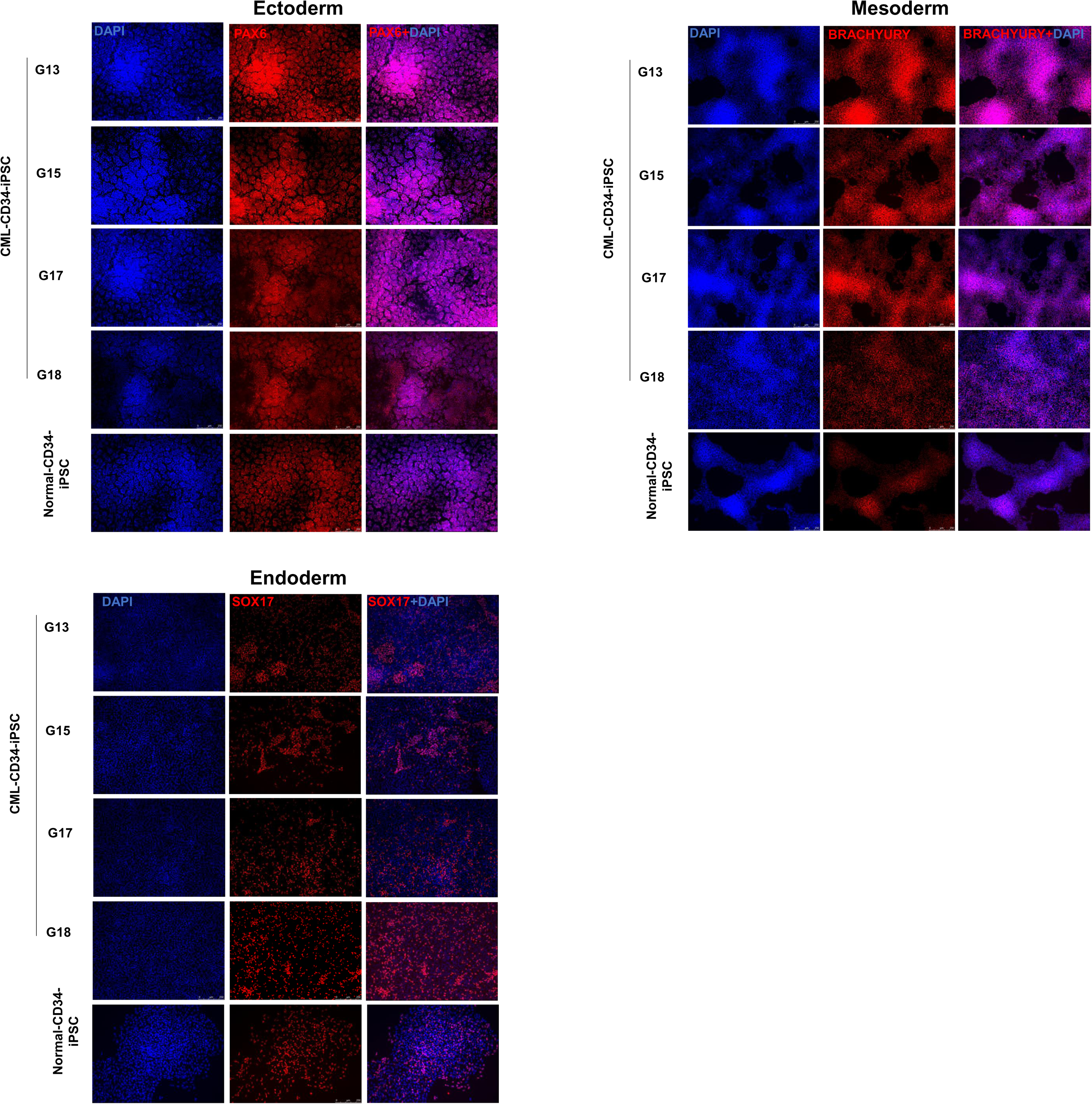

**Figure S4.**
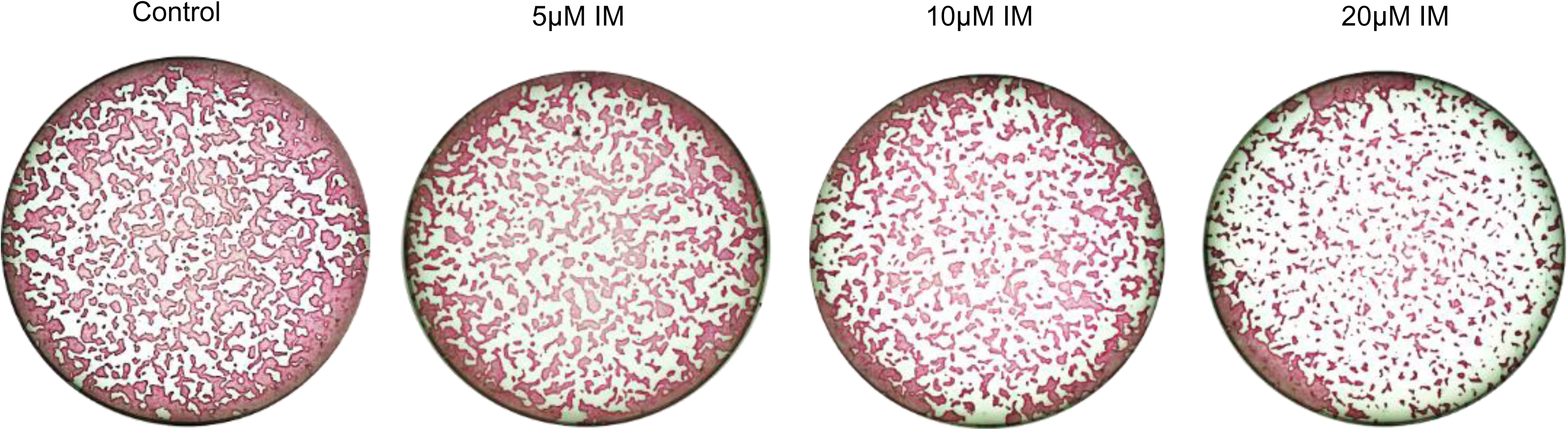

**Figure S5.**
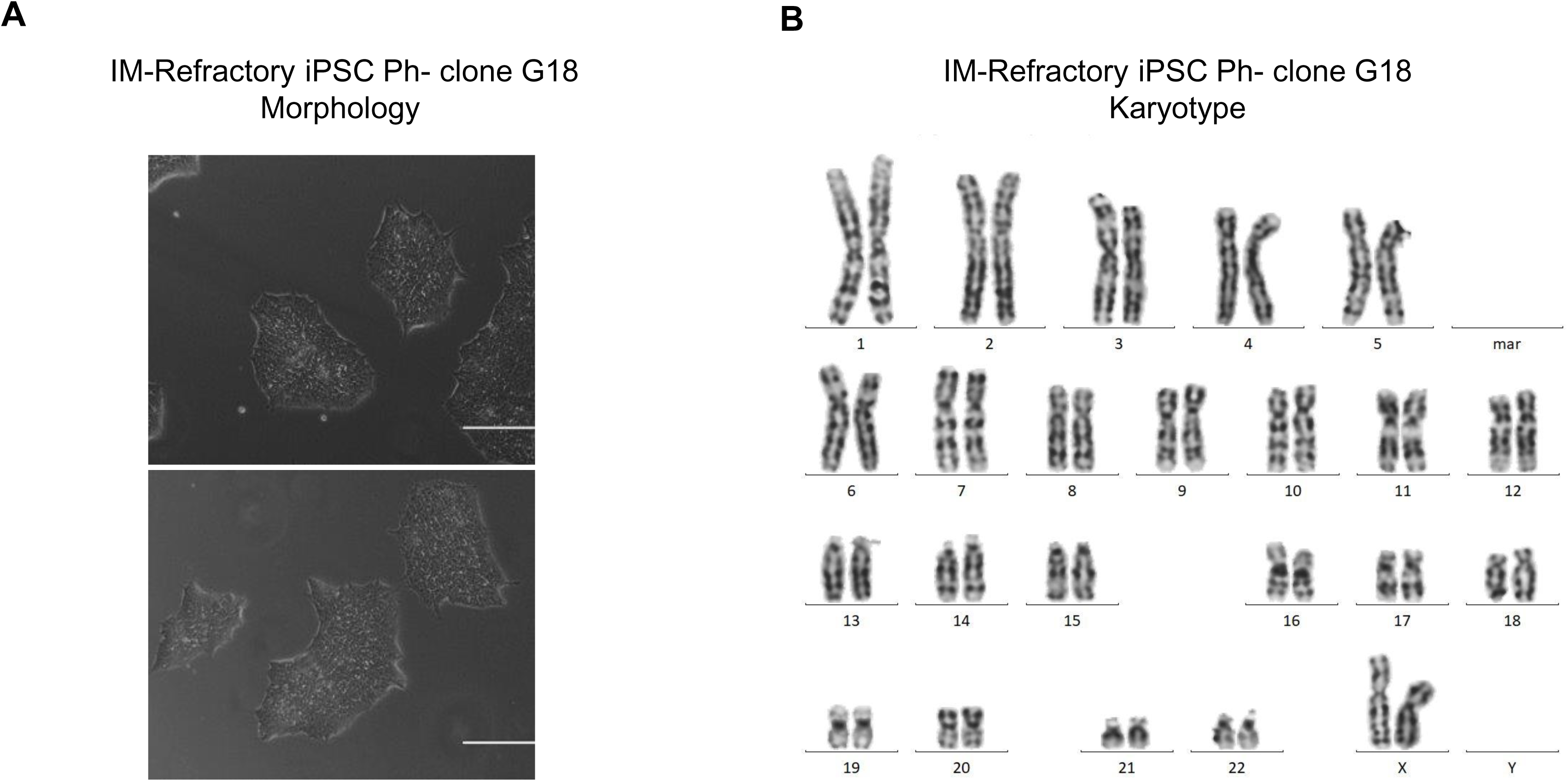

**Figure S6.**
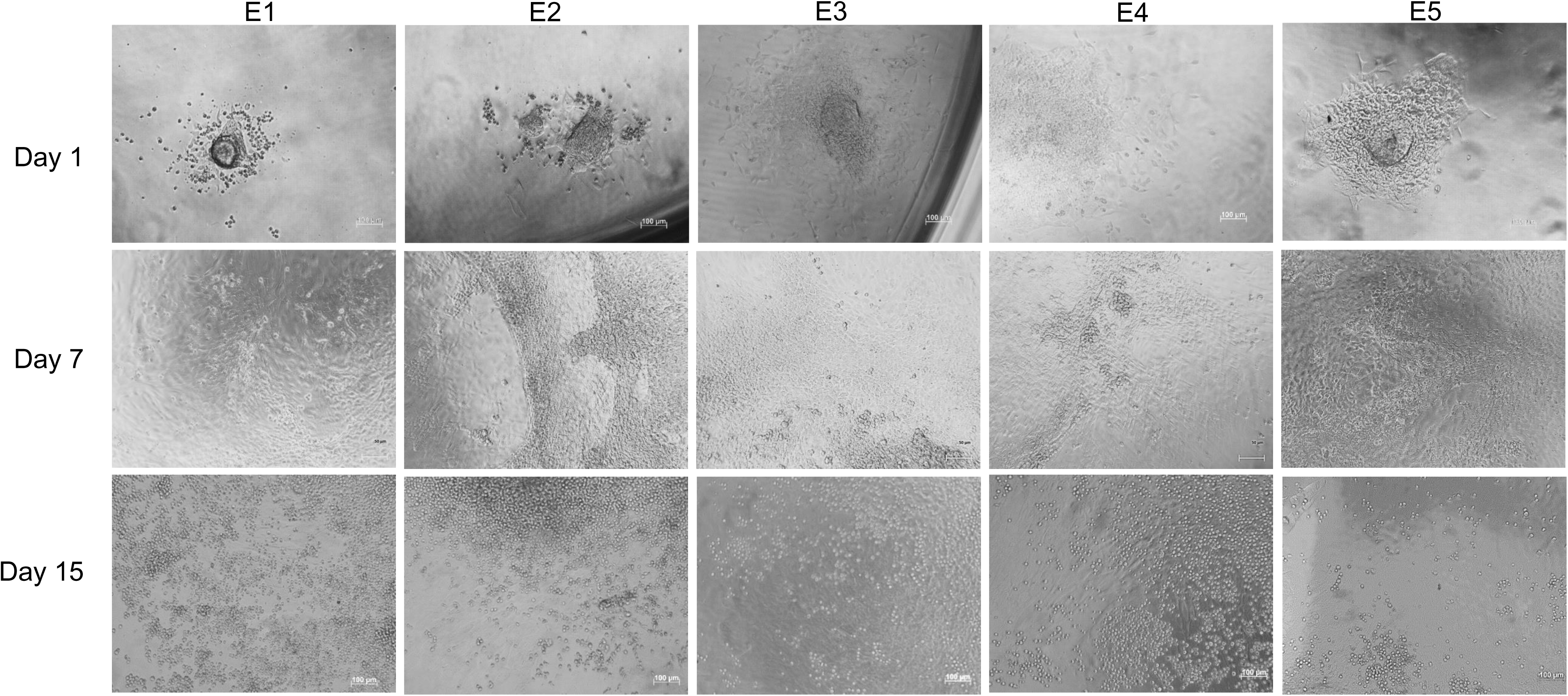

**Table S1.**
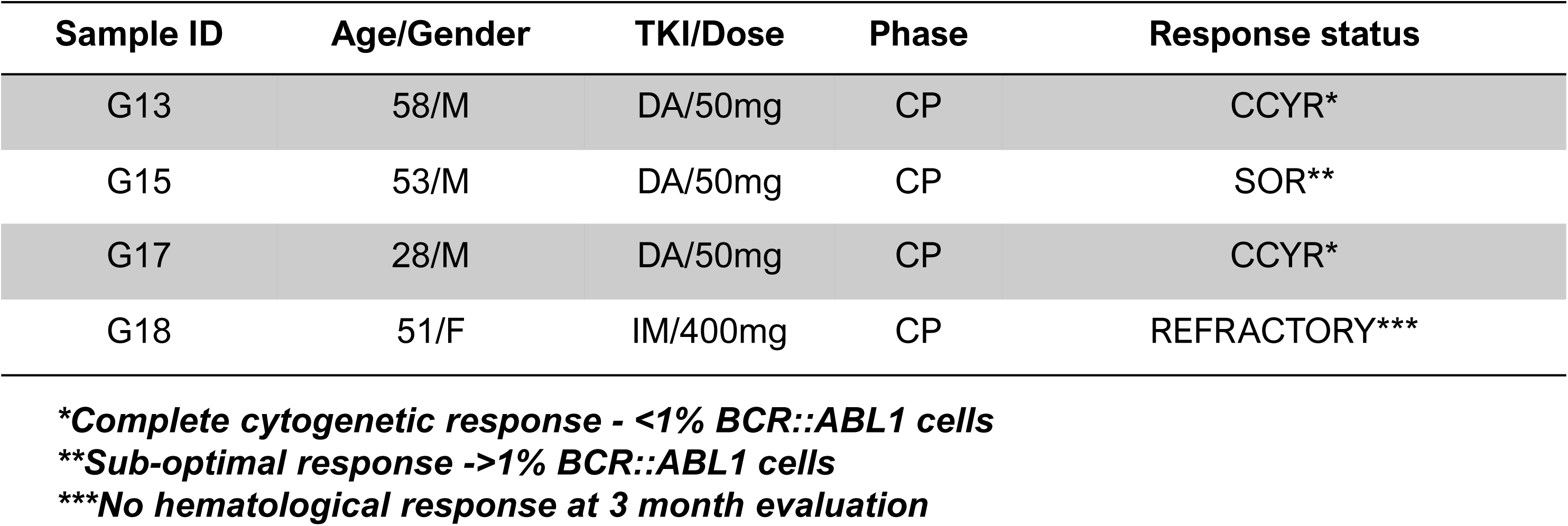

