## Supplemental Data 1 for "Inhibition of BCR::ABL1 tyrosine kinase activity Aids in the Generation of Stable Chronic Myeloid Leukemia Induced Pluripotent Stem Cells"

**Figure S1. Expansion of CD34+ cells.**

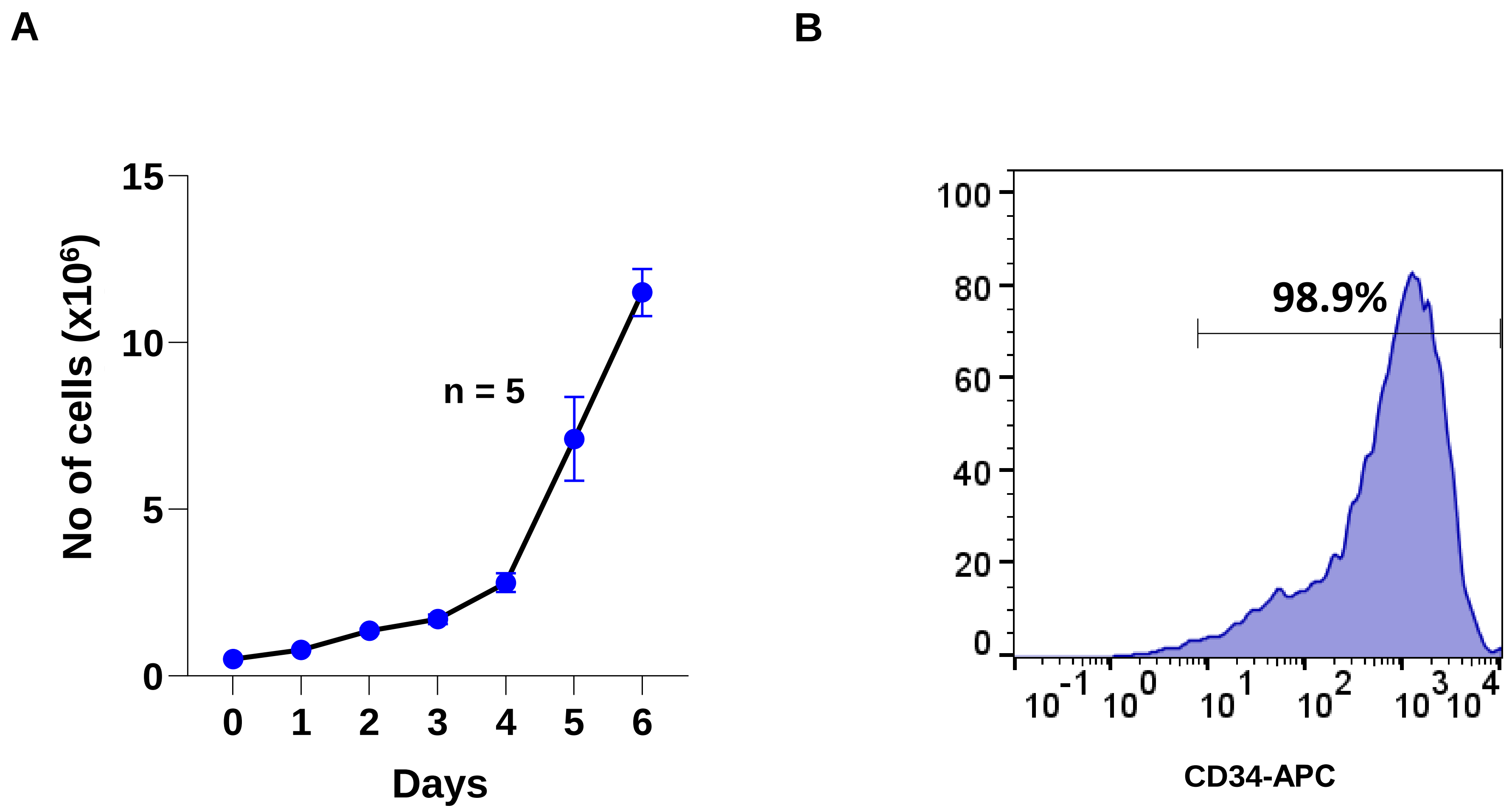


(**A**) Representation of the growth rate of $CD34^{+}$ cultured in the expansion medium. $CD34^{+}$ cells proliferated in the expansion medium till day 6. **(B)** Flow cytometry analysis showing the purity of the $CD34^{+}$ population after expansion. CML $CD34^{+}$ cells showed >90% CD34 Purity.

**Figure S2. Characterization of CML-CD34-iPSC clones.**

**A**

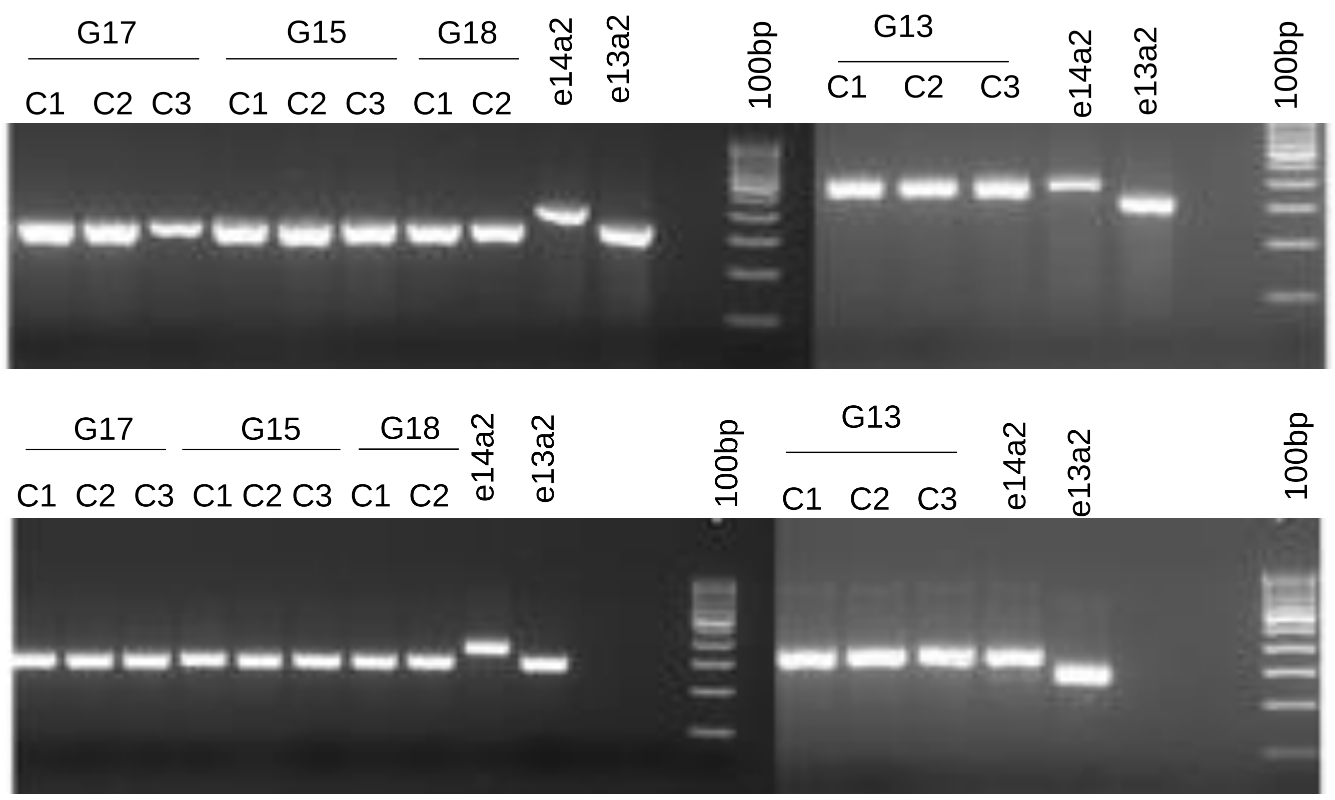


| Patient ID | Transcript Type |
|---|---|
| G17 | e13a2 |
| G15 | e13a2 |
| G18 | e13a2 |
| G13 | e14a2 |

**B**

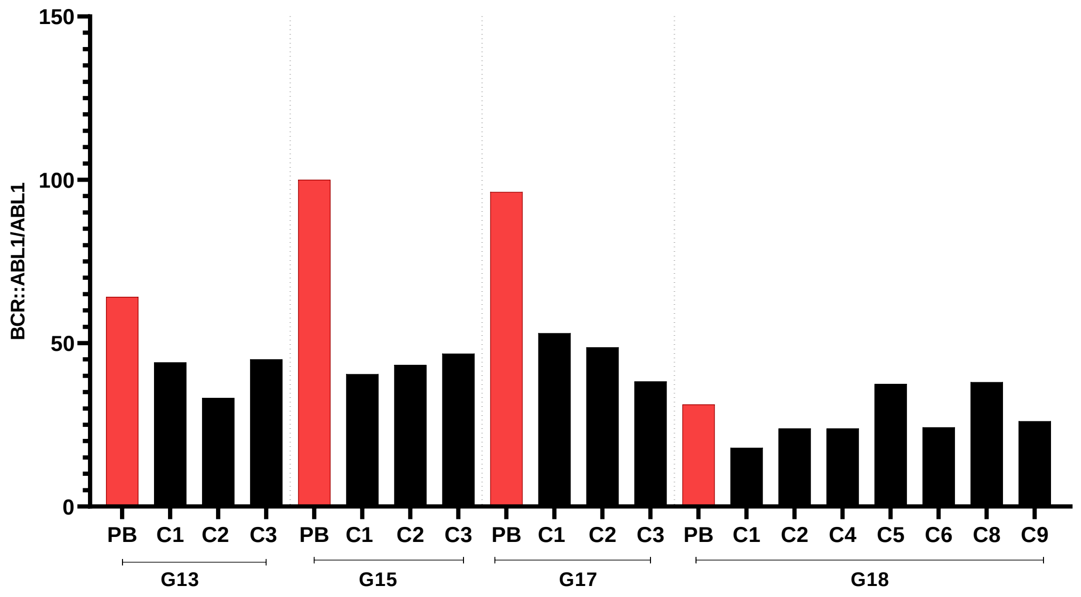


| Patient ID | Clones | Transcript Type |
|---|---|---|
| G17 | C1,C2 and C3 | e13a2 |
| G15 | C1,C2 and C3 | e13a2 |
| G13 | C1,C2 and C3 | e14a2 |
| G18 | C1,C2,C4,C5,C6,C8 and C9 | e13a2 |
| | C7 | Nil |

**C**

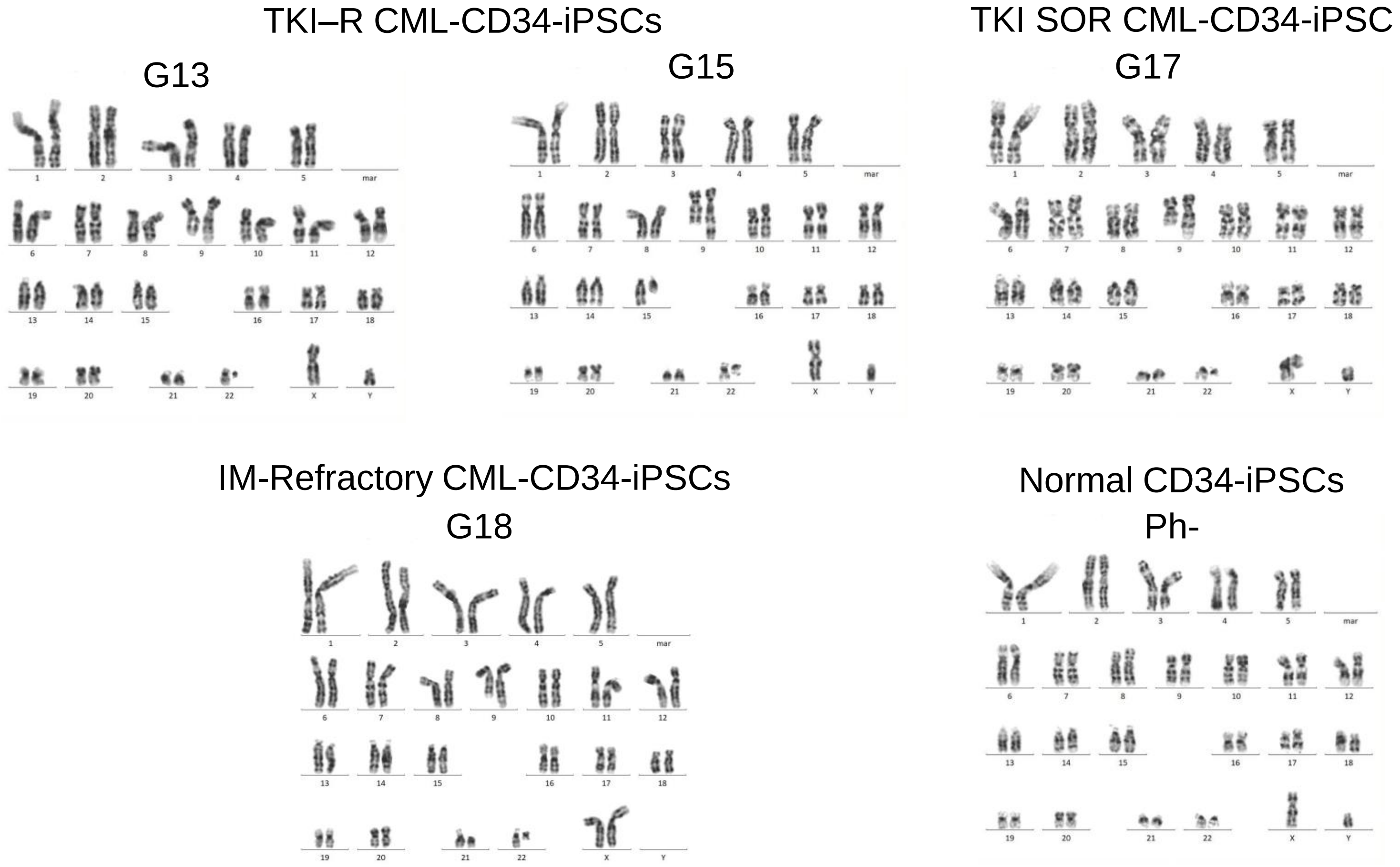


(**A**) Agarose gel image of *BCR::ABL1* transcripts in CML-CD34-iPSC clones. (**B**) Quantification of *BCR::ABL1* transcripts present in the CML-CD34-iPSC clones from 4 patients (normalized to ABL1 expression) by ddPCR. Red bars indicate the *BCR::ABL1* transcripts % in the peripheral blood samples collected at presentation, and the black bars indicate those in the CML-CD34-iPSC clones (cultured with 10μM IM supplementation). (**C)** Karyotyping analysis of iPSC lines showing t(9;22) (q34;q11) translocation in the CML-CD34-iPSC lines except one clone showing normal karyotype. An iPSC line generated from normal-CD34+ cells showing normal karyotype is also shown. #Ph+G13, G17 represents CML-CD34-iPSC clones derived from CML patients, TKI-Tyrosine kinase inhibitor, SOP-Suboptimal response, R-Responder, IM-Imatinib Mesylate, Ph- represent clone showing normal karyotype.

**Figure S3:Trilineage differentiation of CML-iPSC lines and a normal iPSC line.**

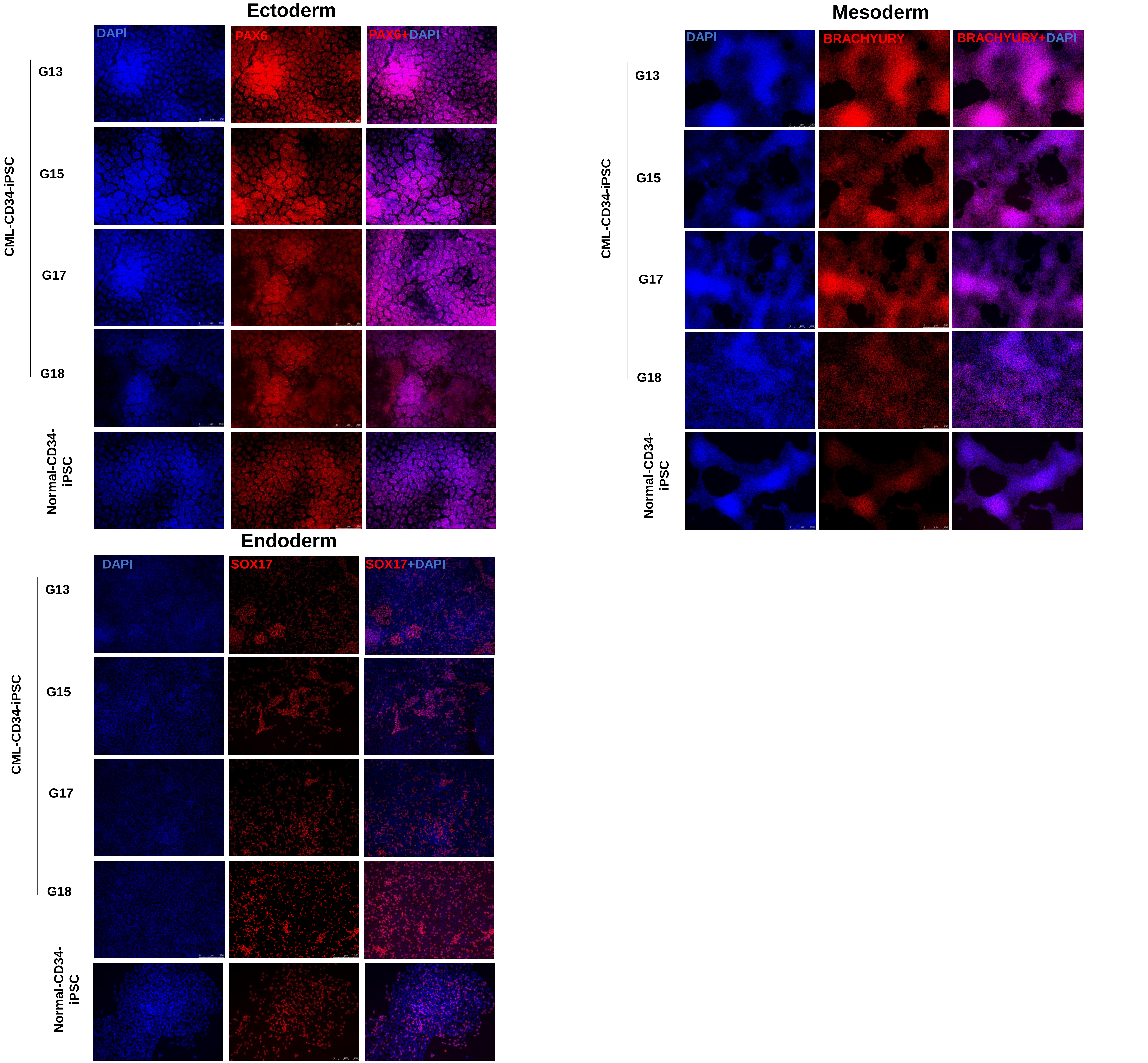


The cells differentiated to ectoderm, endoderm and mesoderm, exhibited the expression of Brachyury, SOX17 and PAX6, as measured by immunofluorescence analysis. DAPI was used for staining the nuclei. G13, G15, G17 and G18 represents CML-CD34-iPSC line. Scale bar:250μM.

**Figure S4; Alkaline phosphatase staining of a healthy donor CD34-iPSC line treated with different concentrations of IM showing a reduction in the colony number after IM treatment.**

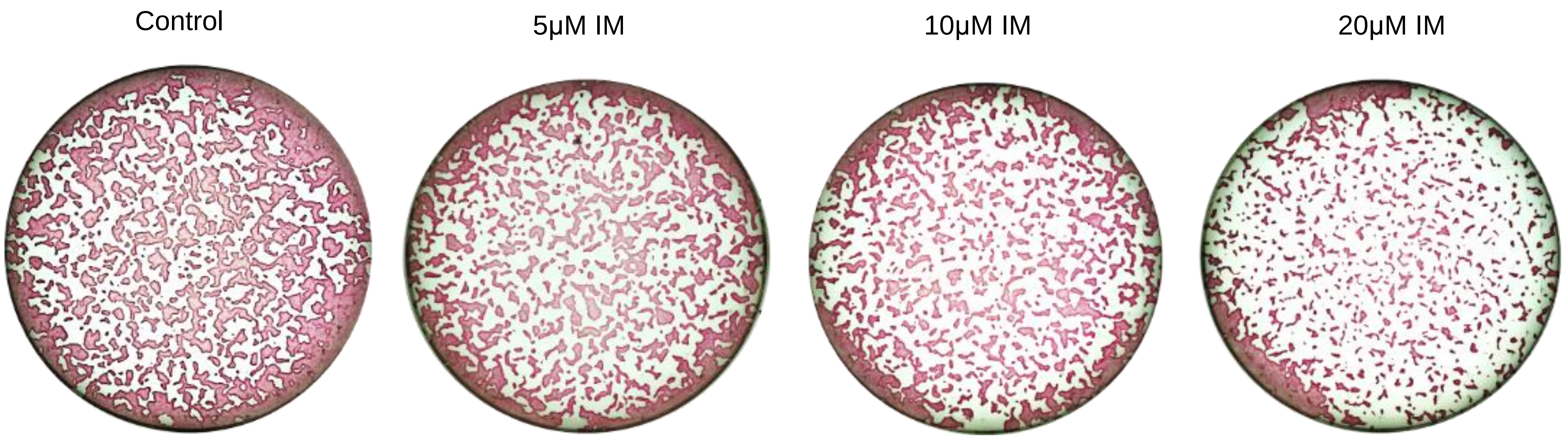

**Figure S5: Reprogramming of IM refractory CML-CD34$^+$ cells.**

**A**

IM-Refractory iPSC Ph- clone G18
Morphology

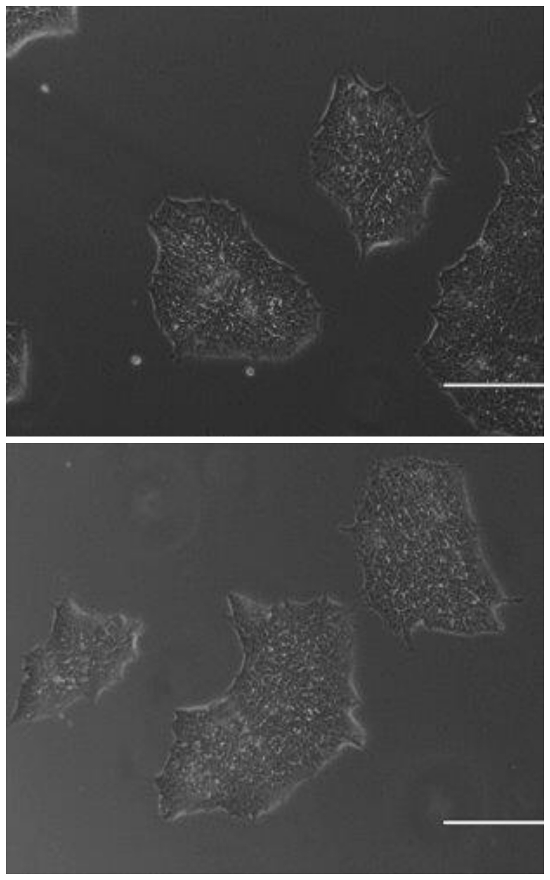

**B**

IM-Refractory iPSC Ph- clone G18
Karyotype

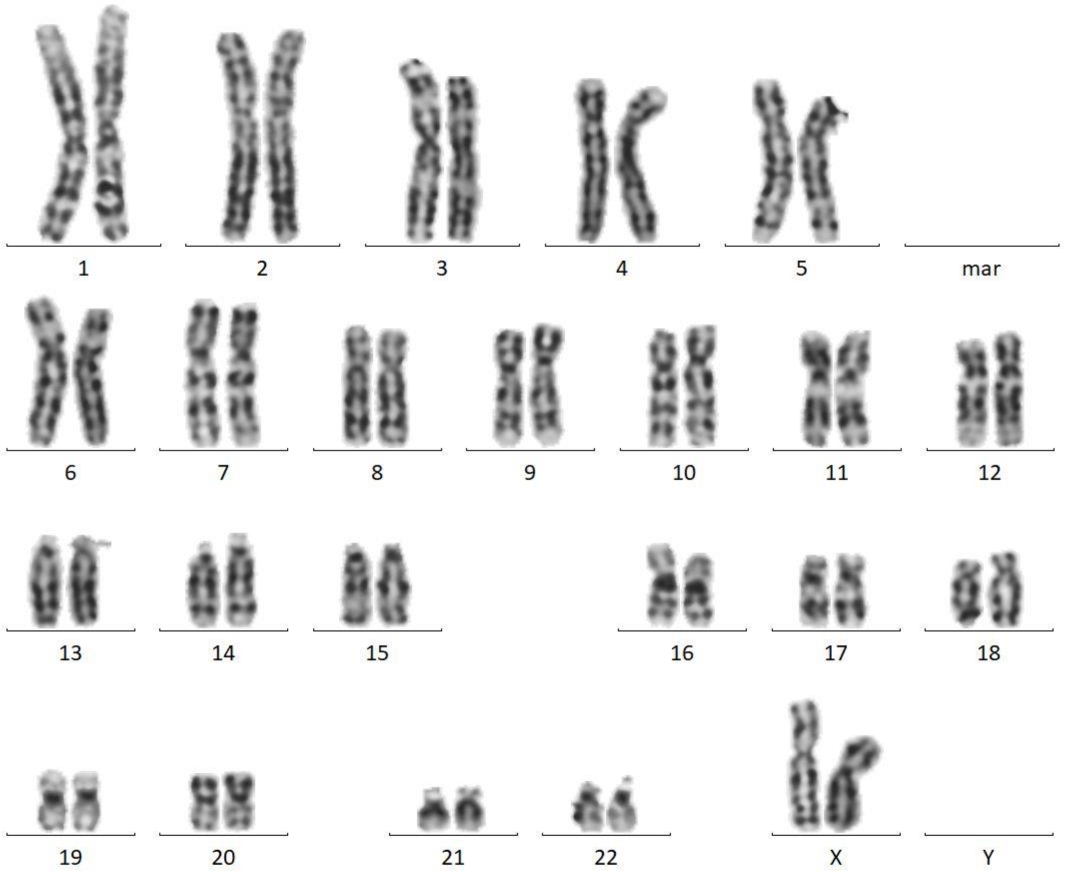


(**A**) Representative images of one Ph-CML-CD34-iPSC colony with typical morphology generated from an IM refractory patient. (**B**) Karyotyping analysis of IM refractory CML-CD34-iPSC clone with typical iPSC morphology showing normal karyotype. IM-Imatinib mesylate, G18- CML patient sample id. Scale bar:50μM.

**Figure S6: Representative images of hematopoietic differentiation of CML-iPSCs.**

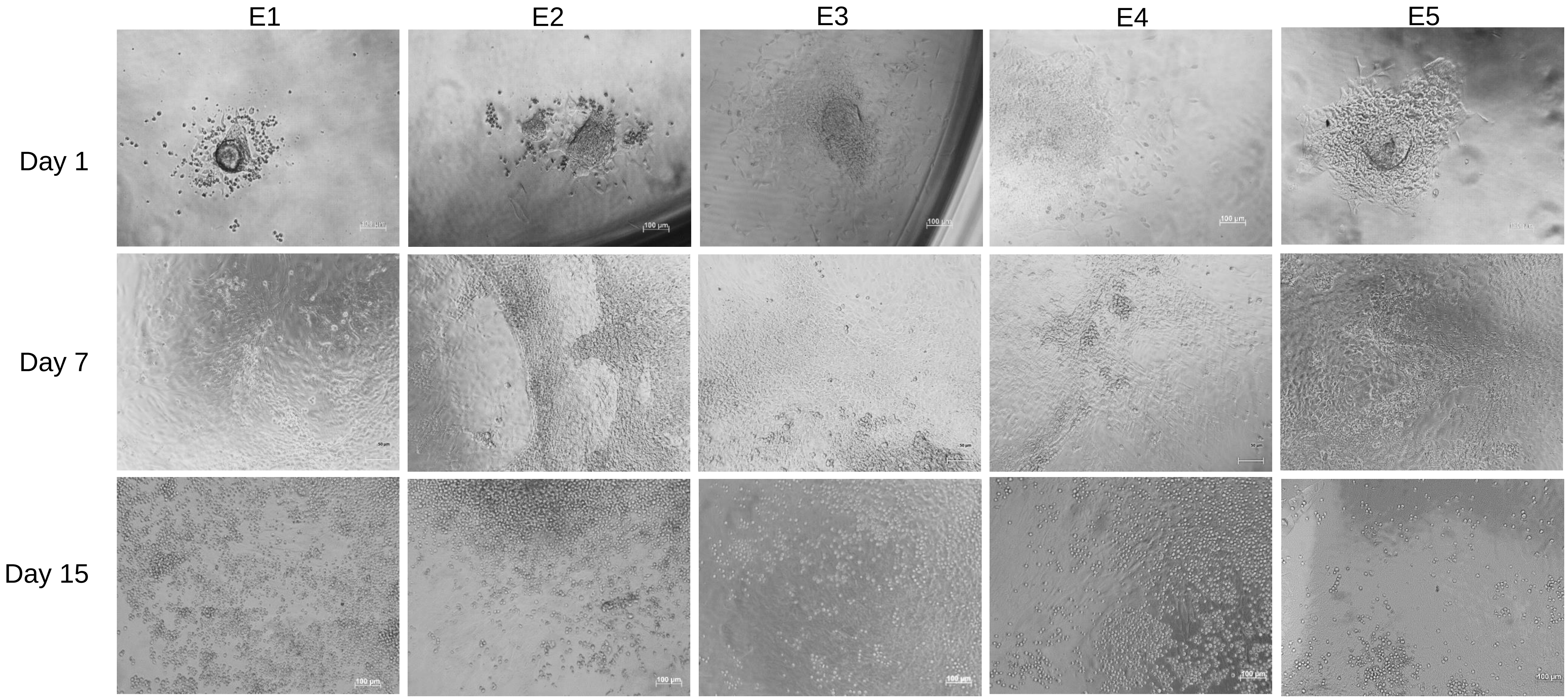

**Table S1: Patient demographics.**

| Sample ID | Age/Gender | TKI/Dose | Phase | Response status |
|---|---|---|---|---|
| G13 | 58/M | DA/50mg | CP | CCYR* |
| G15 | 53/M | DA/50mg | CP | SOR** |
| G17 | 28/M | DA/50mg | CP | CCYR* |
| G18 | 51/F | IM/400mg | CP | REFRACTORY*** |

***Complete cytogenetic response - <1% BCR::ABL1 cells***
*****Sub-optimal response ->1% BCR::ABL1 cells***
******No hematological response at 3 month evaluation***
